# Divergent Cell-Type Specific Hypoxia Responses in Human Stem Cell–Derived and Primary Islets

**DOI:** 10.1101/2025.07.16.665006

**Authors:** Kameron Bradley, Camryn Moore, Matthew Ishahak, Marlie M. Maestas, Daniel A. Veronese-Paniagua, Jeffrey R. Millman

## Abstract

**Background:** The success of stem cell-derived islet (SC-islet) therapy for type 1 diabetes is limited by poor graft survival in the hypoxic post-transplantation microenvironment. While the response of SC-islets to chronic hypoxia has been studied, a direct comparison to primary human islets during the acute hypoxic phase has not been performed. Here, we conduct a comparative single-cell transcriptomic and functional analysis of human SC-islets and primary islets exposed to acute hypoxia (1% O_2_) over 48 hours.

**Results:** Our analysis reveals two divergent response patterns. Primary islets exhibit an energy-conserving response, characterized by a β-cell-specific suppression of identity genes (*PDX1*, *MAFA*) and pro-apoptotic factors like *DDIT3*, alongside a shift toward metabolic quiescence. In contrast, the SC-islet response is characterized by lineage instability, a significant metabolic shift toward glycolysis, and the activation of pro-apoptotic pathways. Functionally, these transcriptomic differences result in a loss of glucose-stimulated insulin secretion in both islet types, but through different mechanisms: a suppression of secretion in primary islets versus dysregulated, glucose-unresponsive insulin release in SC-islets.

**Conclusion:** These findings demonstrate that SC-islets are particularly vulnerable under hypoxic stress, exhibiting an unstable, plastic phenotype. This comparative dataset provides a resource for developing source-specific therapeutic interventions to overcome the hypoxic barrier and improve the efficacy of cell replacement therapies.

## INTRODUCTION

Type 1 diabetes mellitus (T1D) is a chronic metabolic disorder of increasing prevalence, characterized by persistent hyperglycemia due to autoimmune destruction of insulin-producing β cells (1,2). While exogenous insulin injection improves patient outcomes, this therapy does not replicate the precise, dynamic secretion of insulin by endogenous β cells (3). Patients face risks of both hyperglycemia and hypoglycemia, creating significant lifestyle burdens and health risks (4). Primary cadaveric islet transplantation can ameliorate insulin dependence by restoring endogenous insulin secretion and long-term glucose homeostasis (5,6). However, this therapy is limited by the scarcity of donor islets, with estimates suggesting that the supply could only treat ∼0.1% of the global T1D population (7).

To address this limited donor supply, recent strategies have focused on generating stem cell-derived islets (SC-islets) from human pluripotent stem cells (hPSCs) (8–12), including human embryonic stem cells (hESCs) and patient-specific induced pluripotent stem cells (13). These protocols use a multi-stage differentiation that emulates pancreatic organogenesis, progressing from definitive endoderm to pancreatic progenitors and a final endocrine maturation phase (8,9,12,14,15). This process yields three-dimensional aggregates (SC-islets) composed mainly of insulin-producing β-like cells and other endocrine cell types, with sizes similar to primary islets (16–19). These clusters exhibit dynamic, glucose-stimulated insulin secretion (GSIS), demonstrating first-and second-phase insulin release patterns analogous to primary human islets (18). Preclinical success with hPSC derivatives has led to several clinical trials with pancreatic progenitors and SC-islets, with the latter showing early promise in reversing T1D (20,21). This progress supports the potential of SC-islet therapy to address donor islet scarcity, and ongoing clinical trials aim to confirm its safety, durability, and scalability.

Nevertheless, transplanted islets—whether cadaveric or stem cell-derived—are exposed to a hypoxic microenvironment immediately post-engraftment, as a fully functional vasculature does not form until days or weeks post-transplantation (22–27). At many transplant sites, oxygen partial pressures can drop below 15 mmHg (<2% O_2_), leaving islets reliant on diffusion for oxygen supply until new blood vessels form (22,28,29). Insufficient oxygen during this avascular window contributes to the loss of up to 50% of the transplanted β-cell mass, which reduces graft survival and insulin output (30–33). High islet density or larger islet diameters worsen central hypoxia by limiting O_2_ diffusion to the islet core, increasing necrosis and apoptosis (22,23,28,34–36). Hypoxia also blunts GSIS by restricting oxidative phosphorylation and ATP production (37,38), with even moderate hypoxia (3–5% O_2_) suppressing this function (26,39).

Consequently, many recipients require islets from multiple donors to compensate for this early loss of β-cell mass and function.

Both primary and SC-islets have high oxygen demands to fuel insulin secretion, making them vulnerable to hypoxia (17,40–42). In response, islets activate transcriptional programs mediated by hypoxia-inducible factors (HIFs) (43–45). This response shifts metabolism toward glycolysis and promotes angiogenesis to improve survival. However, these adaptations can impair GSIS and lead to the loss of mature β-cell identity (45,46). While some physiological differences in the hypoxic responses of primary and SC-islets are known (31,42), the underlying cell-type-specific transcriptional programs have not been fully characterized.

Here, we investigate the responses of human SC-islets and primary human islets to acute hypoxia *in vitro*. Our comparative analysis reveals source-specific transcriptional programs that drive distinct functional and metabolic outcomes, clarifying the unique vulnerabilities of each islet type for cell replacement therapy.

## RESULTS

### Single-cell transcriptomics identifies distinct cell populations and confirms a conserved core hypoxic stress response across primary and SC-islets

To investigate cell-type-specific responses to hypoxia, we performed single-cell RNA sequencing (scRNA-seq) on SC-islets and primary islets exposed to hypoxia (1% O_2_) for 0, 6, 24, and 48 hours (Fig. 1A). After removal from the incubator, samples were quickly dispersed into single cells and fixed to best preserve their transcriptional state for downstream processing. Fixed samples were hybridized with multiplexing probes to allow for multiplexed scRNA-seq, thereby minimizing batch effects during analysis. Uniform manifold approximation and projection (UMAP) shows effective integration across islet source (Fig. 1B) and time point (Fig. 1C), and graph-based clustering enabled the identification of distinct cell types (Fig. 1D). Cell type identities were validated using established canonical markers (*GCG*+ α, *INS*+ β, *SST*+ δ, *PPY*+ PP, *PRSS1*+ Acinar, *KRT17*+ Ductal, *PECAM1*+ Endothelial, *COL1A1*+ Mesenchymal, *TPH1*+ Enterochromaffin, *TOP2A*+ Proliferating) (Fig. 1E). As expected from previous studies(47,48), we also observed differences in the detected cell types between primary and SC-islets. For instance, primary islets contained several non-endocrine cell types not found in SC-islets, whereas enterochromaffin cells were found only in SC-islets (Fig 1F).

**Figure 1:**
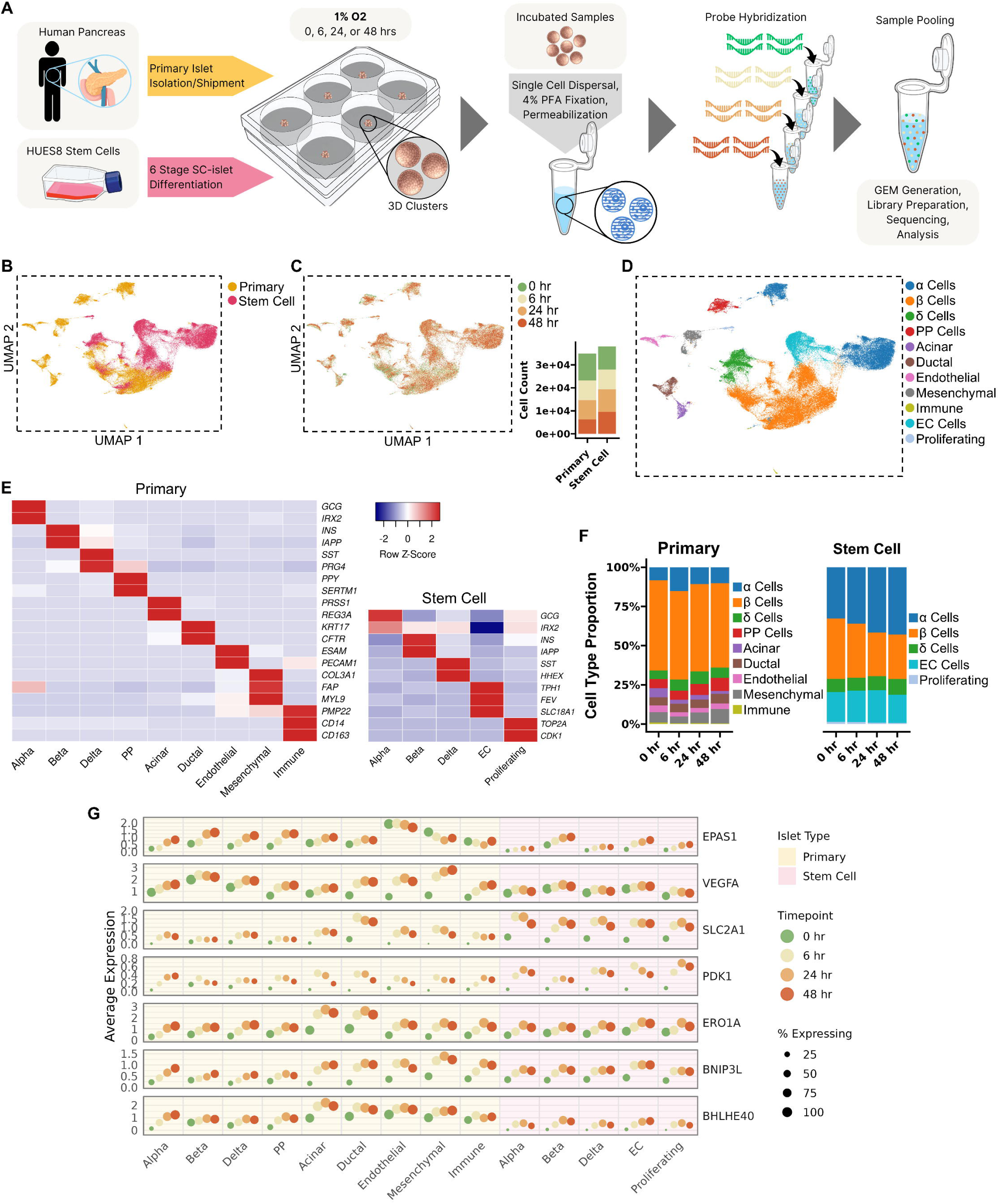
scRNA-seq identifies cell populations and a core hypoxic response in primary and SC-islets. (A) Schematic of the experimental design. Primary human islets and stem cell-derived islets (SC-islets) were exposed to hypoxia (1% O_2_) for 0, 6, 24, or 48 hours. Islet clusters were then dissociated, fixed, and processed for multiplexed single-cell RNA sequencing (scRNA-seq). (B) Uniform Manifold Approximation and Projection (UMAP) of 72,903 integrated cells, colored by islet source (Primary or Stem Cell). (C) UMAP of integrated cells colored by the duration of hypoxia exposure. The inset bar graph shows the number of cells recovered from each source at each timepoint. (D) UMAP of integrated cells colored by annotated cell type identity. (E) Heatmaps validating cell type annotations. The average, scaled expression (Row Z-Score) of canonical marker genes is shown for each identified cell type in primary islets (left) and SC-islets (right). (F) Stacked bar plots showing the relative proportions of each cell type within primary and SC-islet samples across the 48-hour time course. (G) Dot plot illustrating the conserved core hypoxic response. The average expression level (y-axis) and percentage of expressing cells (dot size) are shown for selected canonical hypoxia-response genes across all identified cell types, sources (background color), and timepoints (dot color).

Exposure to acute hypoxia led to a conserved core transcriptional response program across multiple cell types in both primary and SC-islets (Fig. 1G). This involved the upregulation of canonical HIF pathway targets, evidenced by increased expression of the transcription factor *EPAS1* (HIF-2α) itself and its target genes, including oxygen sensors (e.g., *EGLN3* [PHD3]), angiogenic factors (*VEGFA*, *ADM*), and the metabolic regulator *NDRG1*. A concurrent shift towards glycolysis was indicated by increased expression of glucose transporters (*SLC2A1*

[GLUT1], *SLC2A3* [GLUT3]) and key enzymes (*HK2*, *PFKFB3*, *PDK1*). Furthermore, components of the unfolded protein response (UPR) and general stress pathways were broadly induced, including the ER oxidoreductase *ERO1A* and the mitophagy/autophagy-related gene *BNIP3L* (Fig. 1G). Concomitantly, this was paired with a shared downregulation of genes essential for mature endocrine identity and function. This signifies a common strategy of energy conservation via suppression of specialized functions under hypoxic stress, which is detailed further below.

### Hypoxia causes extensive transcriptomic remodeling with source-specific dynamics and increasing deviation over time

We next quantified the magnitude and dynamics of hypoxia-induced transcriptomic remodeling relative to the 0-hour baseline, revealing transcriptomic changes that differed in scale and kinetics between islet sources and cell types (Fig. 2A-B; Supplementary Data 4, 5). During the first 6 hours of hypoxia exposure, SC-α cells showed the largest transcriptional response (Fig. 2A, Supplementary Data 4), with a 14.9-fold greater number of differentially expressed genes (DEGs) compared to primary α cells (5400 vs 363 DEGs), driven primarily by downregulated genes (∼8.9-fold more than upregulated). While the observed number of DEGs progressively increased in primary α cells, peaking at 48 hours (1,484 DEGs), the SC-α cell response was maximal at 6 hours and decreased over time. Both primary and SC-β cells exhibited a large number of DEGs through 24 hours (3,854 and 3,689, respectively). By 48 hours of hypoxia exposure, however, their transcriptional responses diverged: the DEG count in primary β cells further increased to 5,375, driven largely by 4,564 downregulated genes, whereas the DEG count in SC-β cells decreased to 2,989 (Fig. 2A). Diverse transcriptional dynamics were also observed in the other cell types (Supplementary Fig. 1A, Supplementary Data 4, 5).

**Figure 2:**
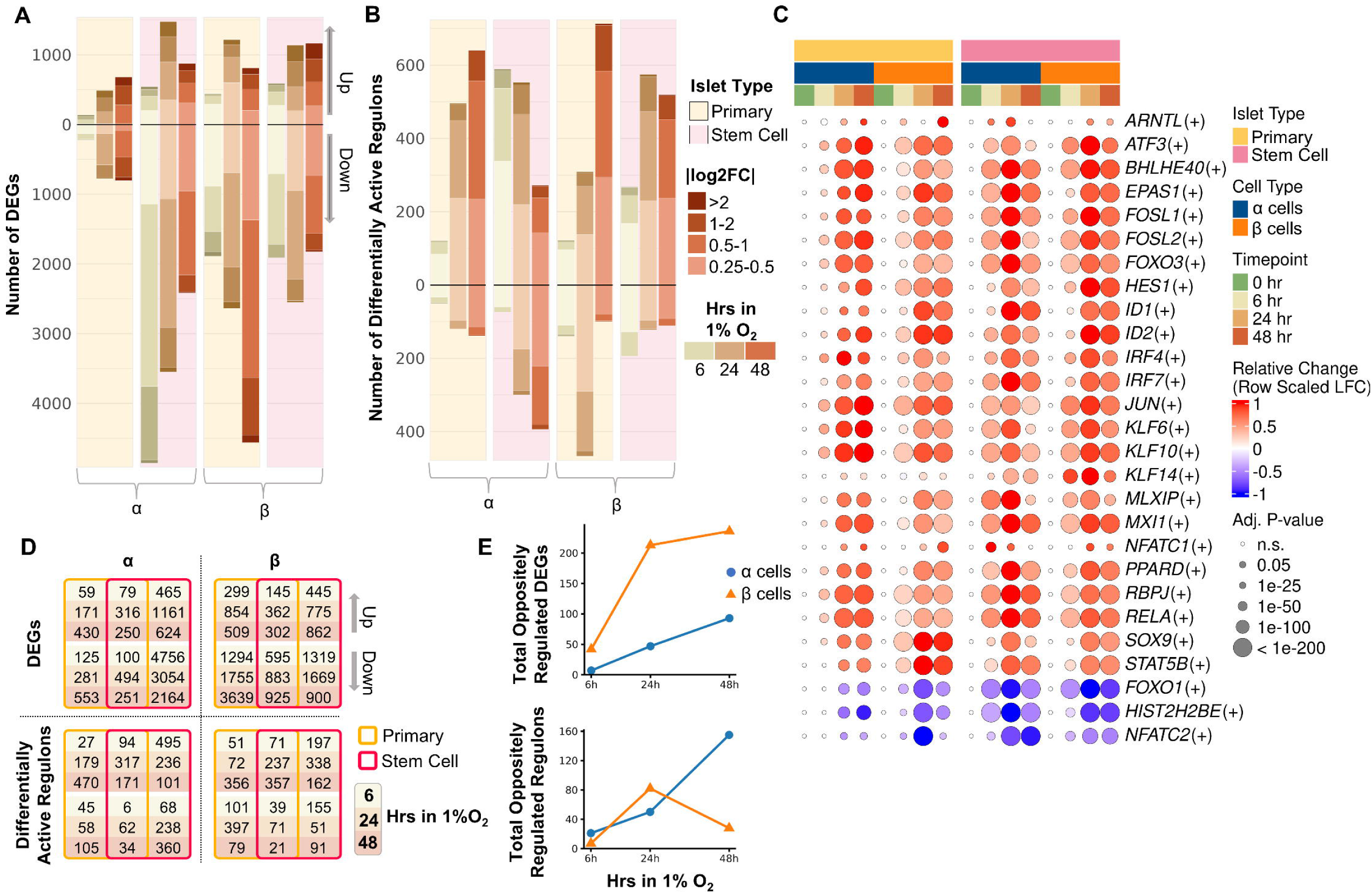
Hypoxia causes extensive transcriptomic remodeling with source-specific dynamics and increasing deviation over time. All comparisons are relative to the 0-hour normoxic baseline for each respective cell type and source. (A) Stacked bar charts quantifying the number of differentially expressed genes (DEGs) in α and β cells from primary and SC-islets at 6, 24, and 48 hours of hypoxia. Bars represent the total number of upregulated (top) and downregulated (bottom) genes. The color indicates the magnitude of the average log_2_ fold-change (|log_2_FC|). (B) Stacked bar charts quantifying the number of differentially active regulons, displayed in the same format as (A). (C) Dot plot showing the activity change for selected regulons that are differentially active in both α and β cells across sources and timepoints. The color represents the relative activity change, scaled to the maximum absolute log-fold change observed for each regulon across all conditions (red = increased activity, blue = decreased activity), and the dot size indicates the adjusted p-value of the change. The top bar indicates islet source and cell type. (D) Matrices summarizing the overlap of DEGs (top) and differentially active regulons (bottom) between primary and SC-islets for α and β cells at 6, 24, and 48 hours. Numbers indicate genes/regulons that are concordantly regulated (up or down in both sources) or uniquely regulated in either primary (yellow) or SC-islets (pink). (E) Line graphs quantifying the total number of genes (top) and regulons (bottom) that are regulated in opposite directions between primary and SC-islets over the 48-hour time course for α and β cells.

To identify the regulatory programs driving these transcriptional shifts, we analyzed the activity within the cellular gene regulatory networks (GRNs) of our samples. This approach infers the activity of a transcription factor by assessing the coordinated expression of its downstream target genes (a “regulon”). Regulons are denoted with a (+) or (-) sign, indicating whether the expression of the transcription factor is positively or negatively correlated with its target gene set, respectively. This analysis of regulon activity changes showed parallel but distinct temporal dynamics (Fig. 2B, Supplementary Data 5). While SC-islet endocrine cells consistently reached their peak number of differentially active regulons at 24 hours (e.g., 853 in SC-α, 697 in SC-β), the response in primary endocrine cells was more prolonged, accumulating and reaching maximal counts at 48 hours (e.g., 813 active regulons in primary β, 780 in primary α).

Direct comparison showed that while core transcriptomic programs were shared, individual responses predominated between primary and SC-islet endocrine lineages (Fig. 2D; Supplementary Data 8). For instance, in β cells at 24 hours, 2,609 DEGs showed changes unique to primary β cells (accounting for ∼68% of total primary DEGs when including oppositely regulated genes), and 2,444 DEGs showed effects unique to SC-β cells (∼66% of total SC DEGs). Conversely, only 1,245 genes were concordantly regulated in both sources. The separation in transcriptomic response profiles was also observed at the GRN level, with source-specific shifts in regulon activity (Supplementary Data 8). At 24 hours, for instance, 469 regulons showed effects unique to primary β cells and 389 showed changes unique to SC-β cells (including 81 oppositely regulated regulons), compared to 308 shared concordantly, highlighting source-specific regulatory programs. The number of genes regulated in opposite directions between sources increased over time (Fig. 2E, Supplementary Data 8). In β cells, this count increased ∼5.6-fold from 42 genes at 6 hours to 236 genes at 48 hours, indicating progressively distinct cellular strategies.

### Gene program analysis reveal source[specific and cell[type[specific hypoxic strategies

To identify the transcriptional programs mediating these hypoxic responses, we first defined a core response program as genes and regulons significantly regulated in the same direction across multiple timepoints in at least two of the three major endocrine cell types (α, β, and δ) in both islet sources. The scale of this shared program is shown in Fig. 3A, with its biological functions detailed in Fig. 3B. The core upregulated program represented a complex adaptive response observed in all endocrine cells. This program included: (1) canonical HIF pathway activation, indicated by the EPAS1(+) (HIF-2α) regulon, its co-factor *CITED2*, the oxygen sensor *EGLN3*, and numerous downstream effectors of angiogenesis (*VEGFA*), vascular tone (*ADM*, *ANGPTL4*), and cellular adaptation (*NDRG1*); (2) a metabolic shift to glycolysis, evident from increased expression of glucose transporters (*SLC2A1*, *SLC2A3*) and key glycolytic enzymes (*ENO1*, *GPI*, *HK2*, *PDK1*, *PFKP*, *PFKFB4*); and (3) coordinated activation of the UPR and integrated stress response, encompassing the oxidoreductase *ERO1A*, ER-associated degradation component *EDEM1*, chaperones (*HSPA6*, *HSPB1*), and the key UPR effector *TRIB3*. In addition to these adaptations, the core program included a signature of extracellular matrix and structural remodeling, with upregulation of *VIM*, *DSP*, *SYNPO*, and *PLOD2*, suggesting a physical restructuring of the islet microenvironment. This response was associated with a large set of shared stress-responsive regulons, including those of the AP-1 family (JUN(+), FOSL2(+)), master stress sensors (ATF3(+), ATF4(+), DDIT3(+)), and key developmental pathways (HES1(+), SOX9(+)).

**Figure 3:**
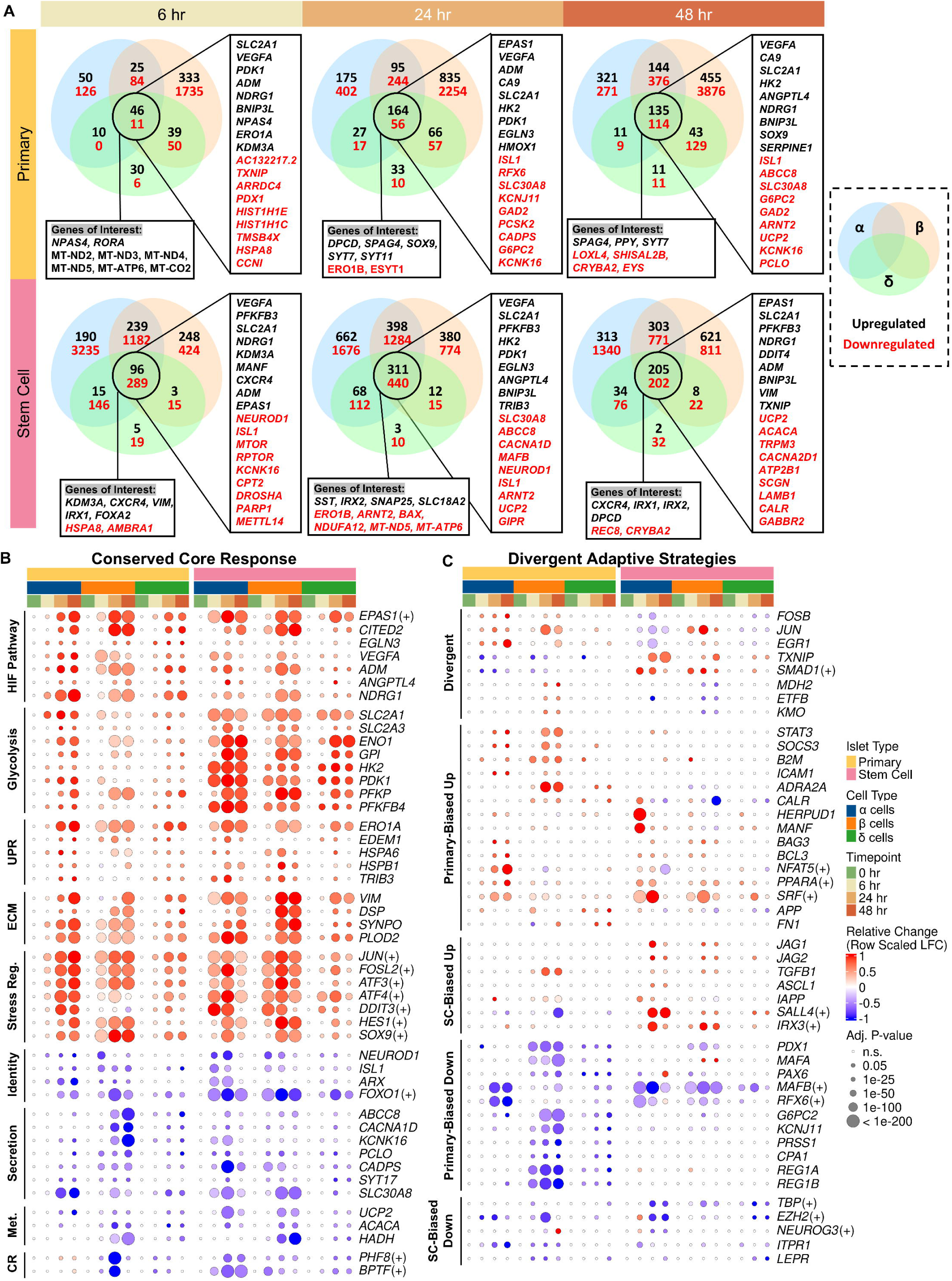
Gene program analysis reveals source-and cell-type-specific hypoxic strategies. (A) Venn diagrams showing the overlap of differentially expressed genes (DEGs) among α (blue), β (orange), and δ (green) endocrine cells at 6, 24, and 48 hours for primary (top row) and SC-islets (bottom row). Numbers indicate upregulated (black) and downregulated (red) DEGs. Boxes highlight representative genes found in the core intersection (regulated in all three cell types). (B) Dot plot visualizing the conserved core response program. Expression changes are shown for representative genes and regulons that are significantly regulated in a similar direction across both primary and SC-islets. Genes are grouped by major biological pathways (HIF Pathway, Glycolysis, UPR, etc.). (C) Dot plot visualizing divergent adaptive strategies. Expression changes are shown for representative genes and regulons that exhibit source-specific regulation. Genes are grouped by their dominant expression pattern (e.g., “Primary-Biased Up,” “SC-Biased Down”). For dot plots (B, C): Dot color represents the relative expression change, scaled to the maximum absolute log-fold change for each gene/regulon across all conditions (red = up, blue = down). Dot size indicates the adjusted p-value. The top bar indicates islet source and cell type. All comparisons are relative to the 0-hour baseline.

Concurrently, the core downregulated program showed a shared pattern of energy conservation through the suppression of mature endocrine identity and function (Fig. 3B). This was driven by the downregulation of transcription factors that define and maintain islet lineages, including *NEUROD1*, *ISL1*, *ARX*, and the pivotal FOXO1(+) regulon. This loss of mature cellular identity markers was coupled with a systematic suppression of the machinery for glucose-sensing and insulin secretion. We observed consistent downregulation of key ion channels (*ABCC8*, *CACNA1D*, *KCNK16*), vesicle-trafficking and docking proteins (*PCLO*, *CADPS*, *SYT17*), and the essential zinc transporter for insulin crystallization (*SLC30A8*).

Furthermore, this metabolic suppression included the downregulation of mitochondrial uncoupling (*UCP2*) and pathways for fatty acid metabolism (*ACACA*, *HADH*), alongside the downregulation of chromatin-modifying regulons (PHF8(+), BPTF(+)). Taken together, this conserved core response involves both the initiation of a broad pro-survival program and the concurrent suppression of energetically expensive, specialized functions.

Beyond the core program, primary islets exhibited a response characterized by suppressed function and promotion of cellular stability (Fig. 3C). This response, shared across the primary endocrine lineages, involved the upregulation of immediate-early transcription factors (*FOSB*, *JUN*, *EGR1*), stress-responsive signaling mediators (*STAT3*, *SOCS3*), and ER chaperones (*CALR*, *HERPUD1*). The islet also initiated a program of intercellular communication and inflammation, evidenced by the upregulation of the major histocompatibility complex component *B2M* and the adhesion molecule *ICAM1*. This program included the specific upregulation of the insulin secretion inhibitor *ADRA2A*, providing a molecular mechanism for functional quiescence. This response was coupled with a coordinated downregulation of genes essential for mature endocrine identity, including the cornerstone transcription factors *PDX1*, *MAFA*, and *PAX6*; glucose-sensing components *G6PC2* and *KCNJ11;* and even several exocrine-associated genes (*PRSS1*, *CPA1*, *REG1A*, *REG1B*), suggesting a broad suppression aimed at preserving energy and maintaining lineage integrity hypoxic stress.

Within this islet-wide strategy, each endocrine cell type made distinct contributions. Primary β cells initiated a complex survival program by simultaneously modifying their metabolism and suppressing cell death pathways. They upregulated genes involved in mitochondrial function (*MDH2*, *ETFB*) and NAD+ synthesis (*KMO*), alongside the β-cell stress-response chaperone *MANF*, indicating an effort to maintain stability. In parallel, primary α cells initiated a specific pro-survival response, activating a suite of anti-apoptotic genes (*BAG3*, *BCL3*) and adaptive regulons (NFAT5(+), PPARA(+), SRF(+)), even as they participated in the broader suppressive theme by downregulating their own identity regulons (ARX(+), MAFB(+), RFX6(+), ISL1(+)). Finally, primary δ cells exhibited a more limited but consistent response profile, upregulating genes involved in cell signaling (*APP*) and extracellular matrix interaction (*FN1*), contributing to the overall adaptive landscape in a targeted manner. Together, these coordinated and cell-type-specific programs depict a strategy of active, controlled quiescence aimed at weathering acute hypoxic stress.

In contrast, SC-islets showed a metabolically active response characterized by lineage instability (Fig. 3C). This difference was evident in the opposing regulation of stress-response programs; while primary islets upregulated immediate-early transcription factors, SC-islets suppressed them (*FOSB*, *JUN*, *EGR1*). Instead of the stabilizing response seen in primary islets, SC-islets upregulated pro-apoptotic programs and the stress sensor TXNIP across endocrine lineages. This unstable state was associated with the re-activation of developmental signaling pathways across multiple lineages, including Notch (*JAG1*, *JAG2*) and TGFβ/BMP (e.g., *TGFB1*, SMAD1(+) regulon), pathways typically suppressed in mature endocrine cells. This instability was also associated with a broad disruption of transcriptional control, evidenced by the shared downregulation of regulons governing core transcriptional machinery (TBP(+)) and chromatin state (EZH2(+), HIST2H2BE(+)).

This widespread instability was driven by distinct, yet coordinated, programs in each cell type. SC-α cells upregulated the neuroendocrine progenitor marker *ASCL1* and ectopically expressed *IAPP*, a peptide typically restricted to β cells. This occurred alongside the specific suppression of the master endocrine progenitor regulon NEUROG3(+), suggesting a shift towards an unstable cell state. SC-β cells also exhibited signs of this maladaptive plasticity. They initiated a strong shift toward glycolysis and displayed lineage infidelity through the activation of progenitor-linked regulons (SALL4(+), IRX3(+)). SC-δ cells were also drawn into this dysregulated state, with a suppression of genes involved in key signaling pathways (*ITPR1*, *LEPR*). Together, these programs depict a strategy of dynamic plasticity over managed stability, rendering SC-islets more vulnerable to hypoxic stress.

### Primary and SC-**β** cells exhibit varied hypoxic adaptations affecting identity, metabolism, and stress pathways

As noted above, primary β cells demonstrated a sustained loss of mature identity markers under hypoxia (Fig. 4A), evidenced by the progressive downregulation of β cell transcription factors including *PDX1*, *MAFA*, *NKX6-1*, *PAX6*, *ISL1*, *NEUROD1*, and *GLIS3*. In contrast, SC-β cells exhibited signs of lineage instability (Fig. 4A). Alongside an initial downregulation of *PDX1*, *PAX6*, and *ISL1*, SC-β cells ectopically upregulated the α-cell hormone *GCG*, the α-associated transcription factors *IRX1* and *IRX2*, and the δ-cell marker *HHEX*. This occurred alongside a paradoxical upregulation of the β-cell maturation factor *MAFA*. Additionally, SC-β cells activated a suite of developmental and progenitor-linked regulons, including SOX1(+), SOX21(+), SOX7(+), POU5F1B(+), SALL4(+), HOXA4(+), HOXC5(+), and PAX8(+).

**Figure 4:**
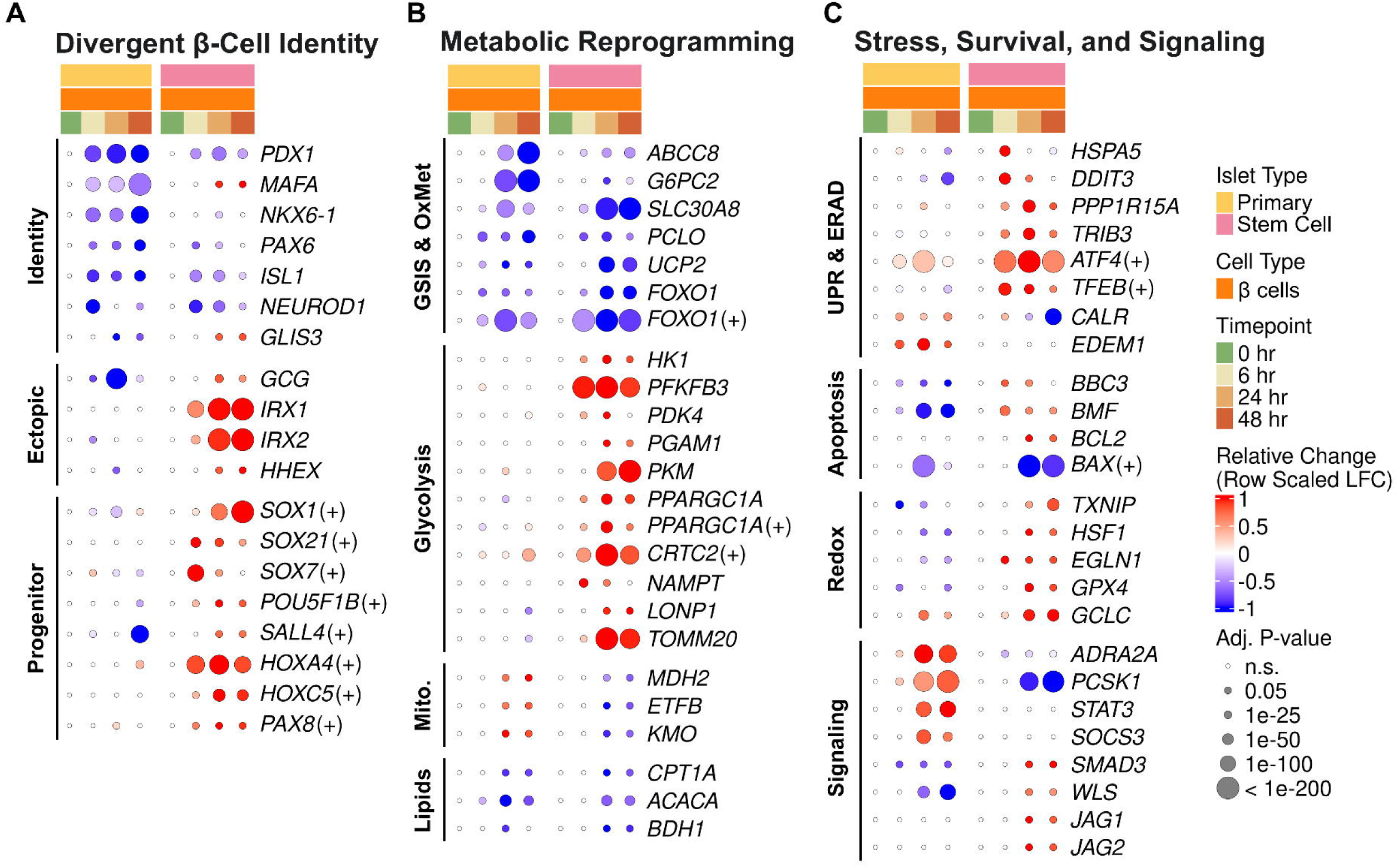
Primary and SC-**β** cells exhibit varied hypoxic adaptations affecting identity, metabolism, and stress pathways. Dot plots comparing gene and regulon expression changes in primary vs. SC-β cells over the 48-hour hypoxic time course. All comparisons are relative to the 0-hour baseline. (A) Divergent β-Cell Identity: Dot plot showing divergent regulation of genes and regulons related to β-cell identity. Genes are grouped by their role in establishing mature identity, their ectopic expression (e.g., non-β-cell hormones), or their association with a progenitor-like state. (B) Metabolic Reprogramming: Dot plot highlighting distinct metabolic reprogramming pathways. Genes and regulons are grouped by function, including glucose-stimulated insulin secretion (GSIS) and oxidative metabolism (OxMet), glycolysis, mitochondrial function (Mito.), and lipid metabolism. (C) Stress, Survival, and Signaling: Dot plot comparing the engagement of stress, survival, and signaling pathways. Genes are grouped by their involvement in the unfolded protein response (UPR) and ER-associated degradation (ERAD), apoptosis, redox balance, and key secretion and signaling pathways. For all panels: Dot color represents the relative expression change, scaled to the maximum absolute log-fold change for each gene/regulon across all conditions (red = up, blue = down). Dot size indicates the adjusted p-value.

Metabolic reprogramming also differed between the two β-cell sources (Fig. 4B). Both primary and SC-β cells suppressed core components of mature GSIS machinery and oxidative metabolism, including sustained downregulation of *ABCC8* (SUR1), *G6PC2*, *SLC30A8* (ZnT8), *PCLO*, *UCP2*, and the *FOXO1* gene and regulon. However, their cellular adjustments differed markedly. SC-β cells exhibited a more pronounced shift towards glycolysis, with sustained upregulation of key enzymes *HK1*, *PFKFB3*, *PDK4*, *PGAM1*, and *PKM*, alongside activation of the primary regulator of mitochondrial biogenesis, *PPARGC1A* (both gene and regulon), and the CRTC2(+) regulon. Further differences were observed in mitochondrial and NAD^+^ metabolism, where primary β cells consistently upregulated *MDH2*, *ETFB*, and the NAD+ synthesis gene *KMO*, while SC-β cells downregulated these same genes. SC-β cells instead upregulated the NAD+ salvage enzyme *NAMPT* and mitochondrial stress/import genes *LONP1* and *TOMM20*. Suppression of fatty acid oxidation (e.g., *CPT1A*, *ACACA*) was evident in both, but SC-β cells more distinctly downregulated *BDH1*, indicating reduced ketone body utilization.

Stress response pathways diverged sharply as well (Fig. 4C). Primary β cells executed a controlled survival program focused on stability and energy conservation. They actively suppressed pro-apoptotic signals, downregulating *DDIT3*, *BBC3* (PUMA), and *BMF*, while bolstering protective ER-associated degradation (upregulating *CALR* and *EDEM1*). This was coupled with a managed redox state, characterized by the downregulation of the stress sensor *TXNIP* and heat shock activator *HSF1* alongside an increase in glutathione synthesis (*GCLC*). Primary β cells appeared to trend toward functional quiescence while maintaining their insulin processing capacity, upregulating both the secretion inhibitor *ADRA2A* and the proinsulin processing enzyme *PCSK1*, while also activating JAK-STAT signaling components (*STAT3*, *SOCS3*).

Conversely, SC-β cells had a more aberrant response. They initiated a more robust ER stress/UPR, evidenced by upregulation of *HSPA5*, *PPP1R15A*, *TRIB3*, and the key pro-apoptotic factor *DDIT3*, along with activation of the ATF4(+) and TFEB(+) regulons. This pro-apoptotic signaling was reinforced by the induction of *BBC3* and *BMF*, yet occurred despite a seemingly contradictory upregulation of the anti-apoptotic *BCL2* and downregulation of the BAX(+) regulon, suggesting a complex and unstable apoptotic balance. Their hyperactive redox state was marked by upregulation of *TXNIP*, *GPX4*, and *HSF1*. Functionally, rather than preserving mature machinery, they downregulated the inhibitory *ADRA2A* and the essential processing enzyme *PCSK1* while broadly reactivating developmental signaling pathways including Notch (*JAG1*, *JAG2*), TGFβ (*SMAD3*), and Wnt (*WLS*). This divergence was also reflected in the opposing regulation of the core HIF pathway inhibitor *EGLN1* (PHD2), which was consistently upregulated in SC-β cells but showed late downregulation in their primary counterparts.

### Immunohistochemistry reveals spatial dynamics of the hypoxic response and validates loss of **β**-cell identity

To validate our transcriptomic findings at the protein level and understand their spatial organization within the three-dimensional islet structure, we performed immunohistochemistry at each hypoxic timepoint (Fig. 5A, F). Consistent with the transcriptional downregulation of key identity genes, we observed a progressive loss of *PDX1* protein in both primary and SC-islets. By 24 and 48 hours, *PDX1* staining was largely restricted to the nuclei of cells in the islet periphery, while being absent from the core (Fig. 5A, F). This spatial pattern provides direct evidence that the most severely hypoxic cells at the islet center lose a critical marker of mature β-cell identity.

**Figure 5:**
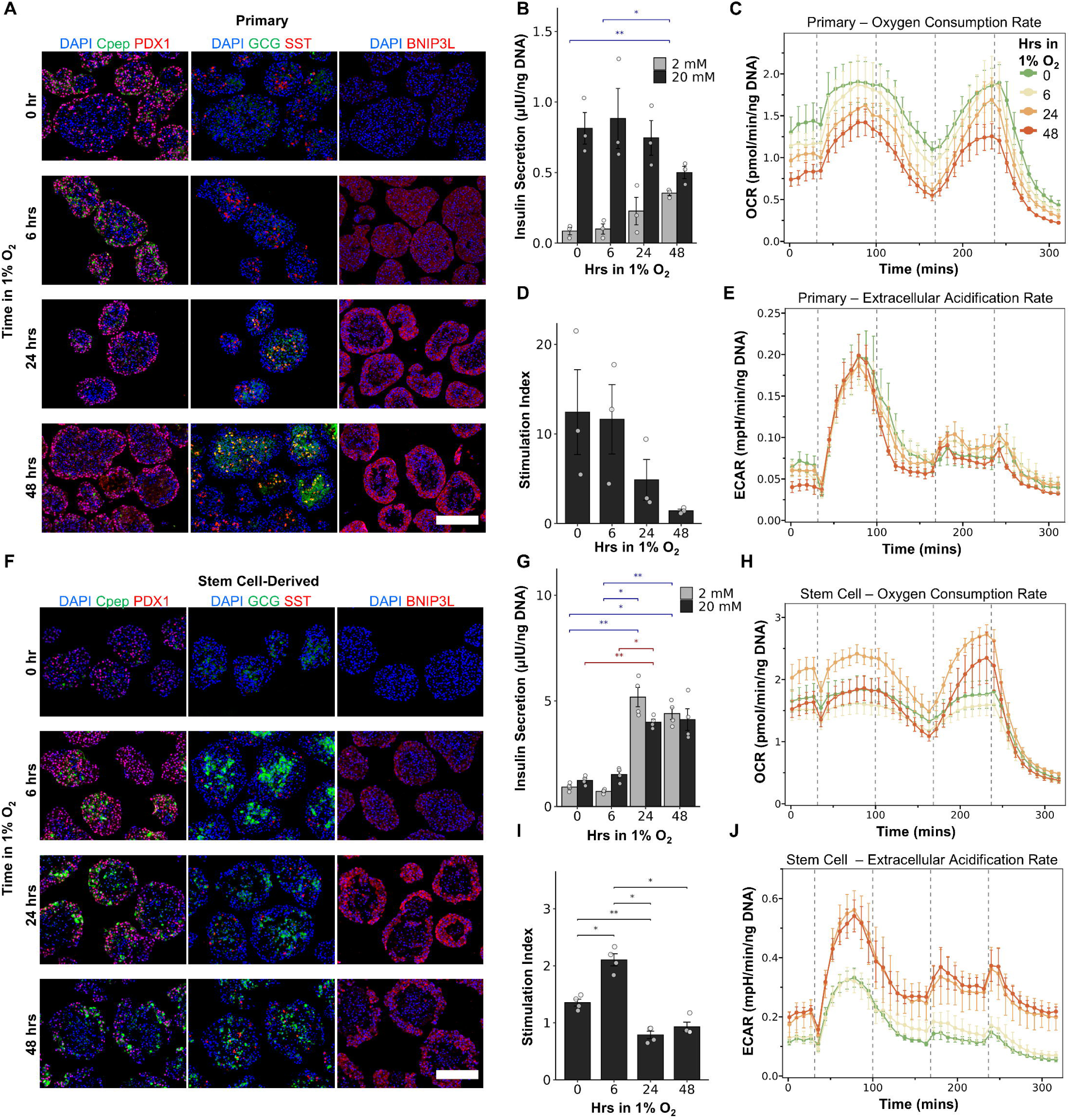
Divergent functional, metabolic, and cellular responses of primary and SC-islets to acute hypoxia. Functional, metabolic, and immunohistochemical analyses of primary islets (top row, A-E) and SC-islets (bottom row, F-J) after exposure to 1% O_2_ for 0, 6, 24, or 48 hours. (A, F) Representative immunohistochemical images of (A) primary and (F) SC-islet sections at each timepoint. Panels show co-staining for DAPI (blue), C-peptide (Cpep, green), and PDX1 (red); DAPI, glucagon (GCG, green), and somatostatin (SST, cyan); and DAPI and BNIP3L (red). Scale bars = 200 µm. (B, G) Absolute insulin secretion in response to low (2 mM, grey bars) and high (20 mM, dark bars) glucose for (B) primary and (G) SC-islets. (C, H) Oxygen consumption rate (OCR) of (C) primary and (H) SC-islets, measured via Seahorse metabolic flux analysis. Different colored lines represent islets recovered after the indicated duration of hypoxia. The traces show metabolic responses to sequential injections (indicated by dashed lines) of glucose (20 mM), oligomycin (1.5 µM), FCCP (0.5 µM), and finally rotenone/antimycin A (0.5 µM). (D, I) Glucose-stimulated insulin secretion (GSIS) stimulation index (SI) for (D) primary and (I) SC-islets, calculated as the ratio of high-to-low glucose secretion. (E, J) Extracellular acidification rate (ECAR) of (E) primary and (J) SC-islets, measured concurrently with OCR. *Data in (B, D, G, I) are presented as mean ± SEM of n=3 (primary) or n=4 (SC-islets) biological replicates; individual data points are shown as open circles. Data in (B, E, G, J) are representative traces from a single experiment, with lines showing the mean ± SEM of technical replicates. For insulin secretion (B, G) and stimulation index (D, I), statistical significance was assessed using repeated measures ANOVA followed by pairwise t-tests with Bonferroni correction for post-hoc analysis. Significance is denoted as *p < 0.05, **p < 0.01.

Furthermore, we examined the expression of the mitophagy-related protein *BNIP3L*, whose transcript was part of the core hypoxic response program. At 6 hours, *BNIP3L* protein was induced throughout both primary and SC-islets, indicating a widespread initial stress response.

However, by 24 and 48 hours, a distinct pattern emerged: strong *BNIP3L* staining was concentrated in the viable cells of the islet periphery, coinciding with the formation of a necrotic core. This suggests that while the entire islet initially senses hypoxia, only the cells in the better-oxygenated periphery are able to mount a sustained protective response, whereas cells in the core succumb to overwhelming stress and cell death.

### Source-specific transcriptomic responses correlate with distinct alterations in islet function and metabolism

Functional assays confirmed that these contrasting transcriptomic programs lead to disparate physiological consequences (Fig. 5). Primary islets exhibited a significant, time-dependent loss of glucose responsiveness (One-way RM ANOVA for SI, p=0.032), with the mean stimulation index (SI) dropping from 12.4 ± 4.7 (n=3) at 0 hours to 1.4 ± 0.2 at 48 hours of hypoxia (Fig. 5D). This loss of function correlated strongly with the transcriptomic downregulation of β-cell identity factors (*PDX1*, *MAFA*, *PAX6*) and the specific upregulation of the secretory inhibitor *ADRA2A* (Fig. 4A, C). SC-islets displayed a transient increase in SI at 6 hours, followed by a complete loss of glucose responsiveness (SI < 1) at 24h and 48h (Fig. 5I). This loss occurred despite markedly increased absolute insulin secretion at later time points (Fig. 5F), indicative of dysregulated, glucose-unresponsive secretion. This is possibly due to a combination of their transcriptional identity instability (Fig. 4A) and the initiation of apoptotic pathways (Fig. 4C), which can compromise membrane integrity and lead to insulin leakage.

Metabolic flux analyses, performed after a 2-hour normoxic recovery period to ensure stable baseline measurements for the assay, revealed corresponding differences in the persistent metabolic state of the islets. Aligning with the suppression of oxidative pathway gene expression (e.g., *UCP2*, *CPT1A*, *IDH1*), primary islets showed a progressive decrease in oxygen consumption rate (OCR) across all respiratory states (Fig. 5C). Notably, despite the transcriptional upregulation of key glycolytic genes (Fig. 3B), their extracellular acidification rate (ECAR) remained relatively stable (Fig. 5E), suggesting a strategy of overall metabolic quiescence rather than a compensatory shift to glycolysis. In contrast, SC-islets maintained OCR through 6 hours but showed elevated maximal respiration at 24 and 48 hours (Fig. 5H). Given that this occurred alongside a complete loss of secretory function, this increased respiration is unlikely to represent improved mitochondrial health. Instead, it may reflect a state of inefficient mitochondrial coupling or altered substrate utilization, a hypothesis consistent with the inferred activation of the PPARGC1A(+) regulon. This model of bioenergetic dysregulation is further supported by a pronounced glycolytic shift, validated by the increased ECAR at 24 and 48 hours (Fig. 5J), which directly reflects the transcriptional upregulation of key glycolytic drivers *PFKFB3* and *PDK4* (Fig. 4B).

## DISCUSSION

The challenge of early post-transplantation hypoxia impedes the success of islet replacement therapies for type 1 diabetes, affecting both primary cadaveric islets and SC-islets. While SC-islets offer a scalable and renewable cell source, their inherent plasticity compared to primary islets requires specific understanding. In this study, we provided a single-cell understanding of islet cell-type-specific responses to hypoxia from both SC-islets and primary islets (Figs. 1-4). Importantly, we identified key functional differences between SC-islets and primary islets in their dynamic response to hypoxia (Fig. 5)

The application of scRNA-seq has provided detailed insights into the biology of both primary and SC-islets, with previous studies characterizing them under basal, diabetic, or other stress conditions (17,47–52). Despite this progress, their response to acute post-transplantation hypoxia remains incompletely understood. While recent work by Wang et al. provided a deep, single-cell characterization of how SC-islets respond to chronic hypoxia (weeks) to identify pro-survival factors (53), a direct comparison with primary islets during the acute phase has not been reported. This study addresses this gap by revealing that while both islet types activate conserved stress pathways, their overall responses diverge. We show that SC-β cells exhibit a plastic and metabolically active response coupled with lineage instability and functional dysregulation. In contrast, primary β cells respond to hypoxia with a strong suppression of their mature identity and a shift towards a quiescent, energy-conserving state. Crucially, these divergent hypoxic trajectories were not only evident at the transcriptomic level but were also consistently reflected in the functional outcomes. The distinct phenotypes of managed functional suppression in primary islets versus dysregulated secretion and metabolic hyperactivity in SC-islets (Fig. 5) were validated using separate biological replicates, lending strong support to the robustness of these source-specific findings. These findings highlight fundamental, source-specific differences relevant for therapeutic development.

These dissimilar hypoxic trajectories likely arise from the inherent differences in developmental maturity and cellular plasticity between the two islet sources. SC-islets, particularly SC-β cells, being products of *in vitro* differentiation, retain greater plasticity than their terminally differentiated adult counterparts (17,47). Our comparative analysis provides strong evidence that this transcriptional flexibility manifests under acute hypoxia as lineage instability in SC-β cells, evidenced by ectopic glucagon (*GCG*) expression and upregulation of α-associated (*IRX1*, *IRX2*) and progenitor-linked (e.g., *HHEX*, SOX TFs) transcripts. This stress-induced instability stands in contrast to the more gradual functional decline previously reported for SC-islets in isolation (53) and was a phenomenon not observed in the primary β cells in our study. Instead, primary β cells executed a strategy of functional retreat, characterized by the progressive and specific downregulation of key identity factors like *PDX1* and *MAFA*.

Our immunohistochemical analysis provides protein-level validation for these divergent transcriptional programs and, importantly, begins to resolve their spatial organization within the hypoxic islet. The observed loss of *PDX1* protein from the islet core in both sources is confirmation of the transcriptional suppression of mature β-cell identity, showing this occurs first in the most severely stressed cells. More strikingly, our analysis revealed a dynamic spatial localization of the mitophagy-related protein *BNIP3L*. While its initial induction was widespread, at later time points its expression was restricted to the viable cells of the islet periphery. This underscores that the divergent transcriptional programs we identified are likely a feature of the surviving cells actively adapting within this oxygen gradient, while the core succumbs to necrosis. This highlights the critical role of the intra-islet oxygen gradient in shaping cellular fate and suggests that future studies using spatial transcriptomics could further resolve these location-specific responses.

Metabolic reprogramming and stress pathway engagement further highlighted these contrasting responses. Primary β cells showed a state of overall metabolic quiescence. This is evidenced by their reduced oxygen consumption and, despite the transcriptional upregulation of glycolytic genes, their stable glycolytic rate, a profile geared entirely towards energy conservation. This metabolic reduction was coupled with the suppression of pro-apoptotic signals like the ER stress factor *DDIT3*. In contrast, SC-β cells had a more metabolically active but unstable response. They attempted to sustain mitochondrial activity via *PPARGC1A* upregulation and initiated a glycolytic shift, validated by their increased ECAR. The metabolic profile of SC-islets under prolonged hypoxia, characterized by a paradoxical increase in maximal oxygen consumption despite a collapse in secretory function, suggests a shift to inefficient mitochondrial coupling. While consuming high levels of oxygen, the cells are unable to efficiently convert this metabolic flux into the ATP required for cellular homeostasis and function. This may result in bioenergetic dysfunction contributing to the observed lineage instability and functional failure. This state of dysregulation was also reflected in the induction of cell death programs, including the upregulation of *DDIT3*, and the opposing regulation of the redox sensor *TXNIP* (upregulated in SC-islets, downregulated in primary islets), which points to a differing, and perhaps more vulnerable, capacity for managing hypoxic stress.

These findings have implications for SC-islet transplantation strategies. The cellular plasticity of SC-islets presents a trade-off during acute hypoxic stress, as it may render the cells vulnerable to the lineage infidelity and functional dysregulation we observed. This is particularly critical because even though SC-islets may establish better long-term vascularization (42), their survival and function during the initial avascular period are essential. Our data show that SC-islets have vulnerabilities distinct from those of primary islets, which requires source-specific protective strategies. This comparative approach complements findings from functional screens that identify general pro-survival factors like *EDN3* (53), by providing a dataset that can be used to identify novel, source-specific therapeutic targets. For instance, interventions for SC-islets could focus on stabilizing β-cell identity (e.g., suppressing GCG expression) or enhancing specific cytoprotective pathways, such as managing ER stress without inducing *DDIT3* or enhancing the specific anti-apoptotic mechanisms involving *BCL2* that we observed. A different strategy for primary islets could involve targeting their upregulation of the secretory inhibitor *ADRA2A*. Finally, further analysis of their metabolic vulnerabilities, such as the opposing NAD^+^ pathways (*KMO* vs. *NAMPT*), could inform future refinements in differentiation protocols or pre-conditioning regimens to enhance hypoxic tolerance.

This study has limitations, including its in vitro nature, the focus on acute hypoxia (up to 48 hours), and the inherent oxygen gradients within 3D islet clusters which cannot be fully resolved by bulk dissociation for scRNA-seq. The scRNA-seq analysis was also based on a single primary islet donor and one SC-islet differentiation batch from a single hESC line (HUES8), which may not capture the full spectrum of inter-donor or inter-line variability.

However, we find it compelling that the distinct adaptive strategies identified for both primary and SC-islets in our transcriptomic data were strongly corroborated by functional, metabolic, and immunohistochemical analyses performed on separate biological replicates (Fig. 5). This consistency across multiple assays and replicates suggests the core divergent cellular programs we report are a robust biological phenomenon. Nevertheless, future work should validate these findings in vivo, explore interventions based on the vulnerable pathways identified (e.g., targeted modulation of metabolic or stress response regulators), and extend analyses to multiple hPSC lines and primary donors.

## CONCLUSION

Our comparative transcriptomic analysis reveals that SC-islets and primary islets have distinct responses to hypoxic stress. Understanding these source-specific cellular responses, particularly the balance between plasticity and stability in SC-β cells, is essential for developing effective approaches to overcome the hypoxic barrier and improve the therapeutic efficacy of SC-islets for type 1 diabetes.

## List of abbreviations

DEG: Differentially Expressed Gene
ECAR: Extracellular Acidification Rate
GRN: Gene Regulatory Network
GSIS: Glucose-Stimulated Insulin Secretion
hESC: Human Embryonic Stem Cell
HIF: Hypoxia-Inducible Factor
hPSC: Human Pluripotent Stem Cell
IHC: Immunohistochemistry
OCR: Oxygen Consumption Rate
SC-islet: Stem Cell-Derived Islet
scRNA-seq: Single-Cell RNA Sequencing
SI: Stimulation Index
T1D: Type 1 Diabetes
UMAP: Uniform Manifold Approximation and Projection
UPR: Unfolded Protein Response

## METHODS

Detailed, step-by-step instructions for most of the methods described here can be found in our previous publication(14). Specific methods are described below.

### Cell lines and culture conditions

The HUES8 human embryonic stem cell (hESC) line was obtained from Dr. Douglas Melton (Harvard University) and maintained in accordance with Washington University Embryonic Stem Cell Research Oversight Committee guidelines (approval no. 15-002). Primary human islets were obtained from Prodo Laboratories with consent from the donor’s relatives for research use. Donor information is provided in Supplementary Data 1. All hESC and human islet experiments were approved by the Washington University Institutional Biological & Chemical (IBC) Safety Committee under protocol number 20889.

Differentiation of hESCs into SC-islets was performed following our established six-stage protocol(14). Briefly, undifferentiated HUES8 hESCs were maintained on Matrigel (Corning; 354277)-coated plates in mTeSR1 medium (StemCell Technologies; 05850) at 37°C and 5% CO_2_. Cells were passaged every 4 days using TrypLE (Gibco; 12-604-013). After 24 hours, medium was replaced daily with mTeSR1 without additional supplements. Differentiation into stem cell-derived islets was performed following an established protocol(14). Cells were seeded at and seeded at 0.63 × 10^6^ cells cm^-2^ in mTeSR1 and 10 μM Y-27632 (Pepro Tech; 129382310MG). After 24 hours, differentiation was initiated using stage-wise media changes and small molecule/growth factor supplementation as outlined in Supplementary Data 1. SC-islets, formed as three-dimensional clusters, were maintained in stage 6 medium(14) and cultured up to 3 weeks, with media changes every 2–3 days.

Primary human islets were sourced as intact clusters from Prodo Laboratories and cultured in CMRL1066 supplemented medium (Corning; 10-060-CV) with 10% heat-inactivated fetal bovine serum (Gibco; 10270106) on an orbital shaker at 115 rpm. Primary islets were used within 5-7 days of arrival.

### Hypoxia exposure

To investigate early to intermediate responses at 0, 6, 24, and 48 hours, SC-islets and primary islets were placed in a standard incubator purged with nitrogen to achieve 1% O_2_ (1% O_2_, 5% CO_2_, balanced N_2_). Both SC-islets (in stage 6 medium) and primary islets (in CMRL + 10% FBS) were cultured in the same manner as at normoxia, with the only difference being the reduced oxygen concentration. Control samples remained at 21% O_2_ in parallel. At each timepoint, islets were harvested for subsequent assays.

### Single-cell RNA sequencing

Single-cell transcriptomic profiling was performed using the Chromium Single Cell Fixed RNA Profiling v1 kit (10x Genomics; 1000414, 1000476) following the manufacturer’s protocol.

Briefly, SC-islets and primary islets exposed to hypoxia for 0, 6, 24, or 48 hours were dissociated into single cells using TrypLE (Gibco; 12604039) for 10 min at 37°C. Samples were fixed and hybridized with multiplexing probes according to the 10x Genomics Fixed RNA Profiling protocols (CG000478, CG000527). Hybridized samples were subsequently transferred to the McDonnell Genome Institute (MGI) at Washington University in St. Louis where reverse transcription, library construction, and sequencing were performed according to the 10x Genomics protocols.

### Single-Cell RNA-seq Analysis

Raw single-cell RNA-seq data were demultiplexed and processed with Cell Ranger (10x Genomics, version 7.0.1) to generate gene-by-cell count matrices aligned to the human reference genome (GRCh38). After initial inspection, all downstream analyses were carried out in R version 4.4.0 using Seurat v5.1.0.

### Data Processing, Integration, and Annotation

Count matrices for each sample were imported using the Read10X function and converted into Seurat objects (CreateSeuratObject). For each sample, low quality cells were filtered out if they showed fewer than 300 features, over 12,500 features, or above 10% mitochondrial content.

Filtered objects were merged into a single Seurat object. Each cell’s RNA assay was converted into the Seurat Assay5 format to facilitate the newer SCTransform pipeline. Normalization and variance stabilization were performed with SCTransform (with vars.to.regress = “percent.mt”), retaining up to 8,000 variable features for downstream analyses. Data were integrated using Harmony with the IntegrateLayers function. Principal component analysis (PCA) was run on the integrated data, retaining 40 principal components for subsequent dimensionality reduction via RunUMAP. Nearest-neighbor graphs (FindNeighbors) and clustering (FindClusters) were performed at a resolution of 0.2. Cell clusters were annotated manually by comparing known marker genes (e.g., INS, GCG, SST, KRT17, CD14) via AverageExpression and cluster-specific marker analysis (FindAllMarkers).

### Differential Expression Analyses

After setting the default assay to “SCT,” each major cell population (e.g., “primary β cells” and “SC-β cells”) was subset and compared across timepoints (0 hr vs. 6 hr, 24 hr, 48 hr). For each comparison, FindMarkers (Wilcoxon test, logfc.threshold = 0.25, min.pct = 0.25, p_val_adj < 0.05) identified differentially expressed genes (DEGs). DEGs were designated as upregulated (positive average log_2_ fold change) or downregulated (negative fold change).

### SCENIC and Regulon Activity

To infer regulon activity, a Loom file of raw counts (SCopeLoomR package) was created for downstream pySCENIC analysis. The resulting regulon activity scores were re-imported as an “AUCell” assay in Seurat. Similar differential analyses (FindMarkers, Wilcoxon test) were performed on regulon activity to identify differentially active gene regulatory networks.

### Glucose-stimulated insulin secretion

GSIS was performed on SC-islets and primary islets following exposure to 1% O_2_ for 0, 6, 24, or 48 hours as previously described(14). Briefly, islets were preincubated in Krebs-Ringer bicarbonate (KREB) buffer containing 2 mM glucose for 1 hour at 37°C, followed by sequential 1-hour incubations in KREB buffer with low (2 mM) and then high (20 mM) glucose. Insulin concentration in the supernatants was measured using a human insulin-specific ELISA (ALPCO; 80-INSHU-E01.1) according to the manufacturer’s instructions. GSIS stimulation index was calculated as the ratio of insulin secreted at high glucose to that at low glucose.

### Metabolic Flux Analysis

OCRs and ECARs were measured using a Seahorse XFe24 Extracellular Flux Analyzer (Agilent). Following exposure to hypoxia for the indicated durations, islets were allowed to recover for approximately 2 hours at normoxia (21% O2) during assay preparation. Seahorse XF24 Islet Capture Microplates (Agilent; 101122-100) were used for the assay. Approximately 70-100 islets per well were seeded into the microplate in Seahorse XF DMEM Medium (Agilent; 103575-100) supplemented with 3 mM glucose, 2 mM glutamine, and 1 mM pyruvate, at pH 7.4. Islets were secured at the bottom of the wells using the provided islet capture screens. The plate was then incubated in a 37°C non-CO2 incubator for one hour prior to starting the assay.

A modified mitochondrial stress test was performed. After establishing a baseline metabolic rate, compounds were sequentially injected through the ports of the sensor cartridge. The injection strategy was as follows: Port A) 20 mM glucose; Port B) 1.5 µM oligomycin (Sigma-Aldrich; O4876); Port C) 0.5 µM FCCP (Sigma-Aldrich; C2920); and Port D) a mix of 0.5 µM rotenone (Sigma-Aldrich; R8875) and 0.5 µM antimycin A (Sigma-Aldrich; A8674) (final concentrations in well). The concentration for the uncoupler FCCP was optimized in prior experiments.

Following the completion of the assay, OCR and ECAR data were normalized to total DNA content per well. Islets were lysed, and total DNA content was quantified using the Quant-iT PicoGreen dsDNA Assay Kit (ThermoFisher; P7589) according to the manufacturer’s instructions.

### Immunohistochemistry (IHC)

For IHC, SC-islets and primary islets were fixed in 4% paraformaldehyde (Electron Microscopy Sciences; 15713) overnight at 4 °C, briefly washed with PBS, and then stored in 70% ethanol at 4 °C until processing (e.g., embedding and sectioning). After deparaffinization, rehydration, and antigen retrieval, slides were permeabilized and blocked in PBS containing 0.1% Triton X-100 and 5% donkey serum for 30 min at room temperature (RT).

Primary and secondary antibodies used to detect C-peptide, PDX1, SST, GCG, and BNIP3L are detailed in Supplementary Data 1, including each reagent’s source, catalog number, working dilution, and other relevant notes. Briefly, slides were incubated with primary antibodies overnight at 4 °C, rinsed in PBS, and then incubated with the appropriate secondary antibodies for 2 h at RT. DAPI (ThermoFisher; D1306) was used for nuclear staining.

Images were acquired on a Leica DMI4000 B and processed using ImageJ. Slides were examined qualitatively to observe expression patterns and morphological features.

## Statistical and Reproducibility

### scRNAseq

For the single-cell RNA sequencing analysis, cluster-level differential expression and regulon activity testing were performed using the Wilcoxon rank-sum test as implemented in Seurat, with p-values adjusted for multiple comparisons using the Bonferroni correction (p < 0.05 as significance threshold).

### Other assays

For functional assays, sample sizes were chosen to ensure adequate statistical power. Specifically, Glucose-Stimulated Insulin Secretion (GSIS) and Stimulation Index (SI) data are from n=3 biological replicates for primary islets and n=4 for SC-islets. Statistical analysis for this data was performed in R (v4.4.0) using the rstatix package. Repeated measures ANOVA was conducted to assess the effects of hypoxia over time, followed by pairwise t-tests with Bonferroni correction for post-hoc comparisons. Key ANOVA assumptions, including normality (via Shapiro-Wilk test) and sphericity (via Mauchly’s test), were evaluated during the analysis.

Seahorse metabolic flux data are presented as a representative experiment, with error bars indicating the standard error of the mean (SEM) of technical replicates. Immunohistochemistry was analyzed qualitatively. Unless otherwise specified, quantitative data are presented as mean ± SEM, and a p-value < 0.05 was considered statistically significant.

## ACKNOWLEDGMENTS

This work was funded by the NIH (R01DK114233, UG3DK142188), Breakthrough T1D (3-SRA-2023-1295-S-B, 2-SRA-2023-1314-S-B), the Edward J. Mallinckrodt Foundation, startup funds from the Washington University School of Medicine Department of Medicine, and the Anita Palmer Corbin Trust to J.R.M. Further support was provided by the Washington University Diabetes Research Center (P30DK020579), particularly for the Seahorse assays. We thank the Genome Technology Access Center at the McDonnell Genome Institute at Washington University School of Medicine for help with genomic analysis. The Center is partially supported by NCI Cancer Center support Grant P30CA91842 to the Siteman Cancer Center from the National Center for Research Resources (NCRR), a component of the National Institutes of Health (NIH), and NIH Roadmap for Medical Research. This publication is solely the responsibility of the authors and does not necessarily represent the official view of NCRR, NIH, nor any other funder. K.B. was supported by Bill & Melinda Gates Foundation through the Gates Millennium Scholars Program. C.M. was supported by the Washington University MARC U-STAR Program (NIH T34GM083914). M.I was supported by Rita Levi-Montalcini Postdoctoral Fellowship in Regenerative Medicine and the NIH (T32DK007120). M.M.M. was supported by the Cellular and Molecular Biology Training grant (T32GM139774). D.A.V.P. was supported by the NSF Graduate Research Fellowship Program (DGE-2139839 and DGE-1745038). We would also like to thank Dr. Chandler Est (Washington University) for early insights into this project and Erika Brown (Washington University) for helpful feedback on the manuscript.

## AUTHOR CONTRIBUTIONS

K.B. and J.R.M. designed all experiments and wrote the manuscript. K.B. and C.M. conducted the wet lab work. K.B., M.I., M.M.M., D.A.V.P. provided key contributions to the sequencing analysis and K.B. conducted this analysis. K.B. and J.R.M. wrote the manuscript. All authors read, had the opportunity to provide feedback, and agreed to the manuscript.

## Supplemental Figure Legend

**Supplementary Figure 1:**
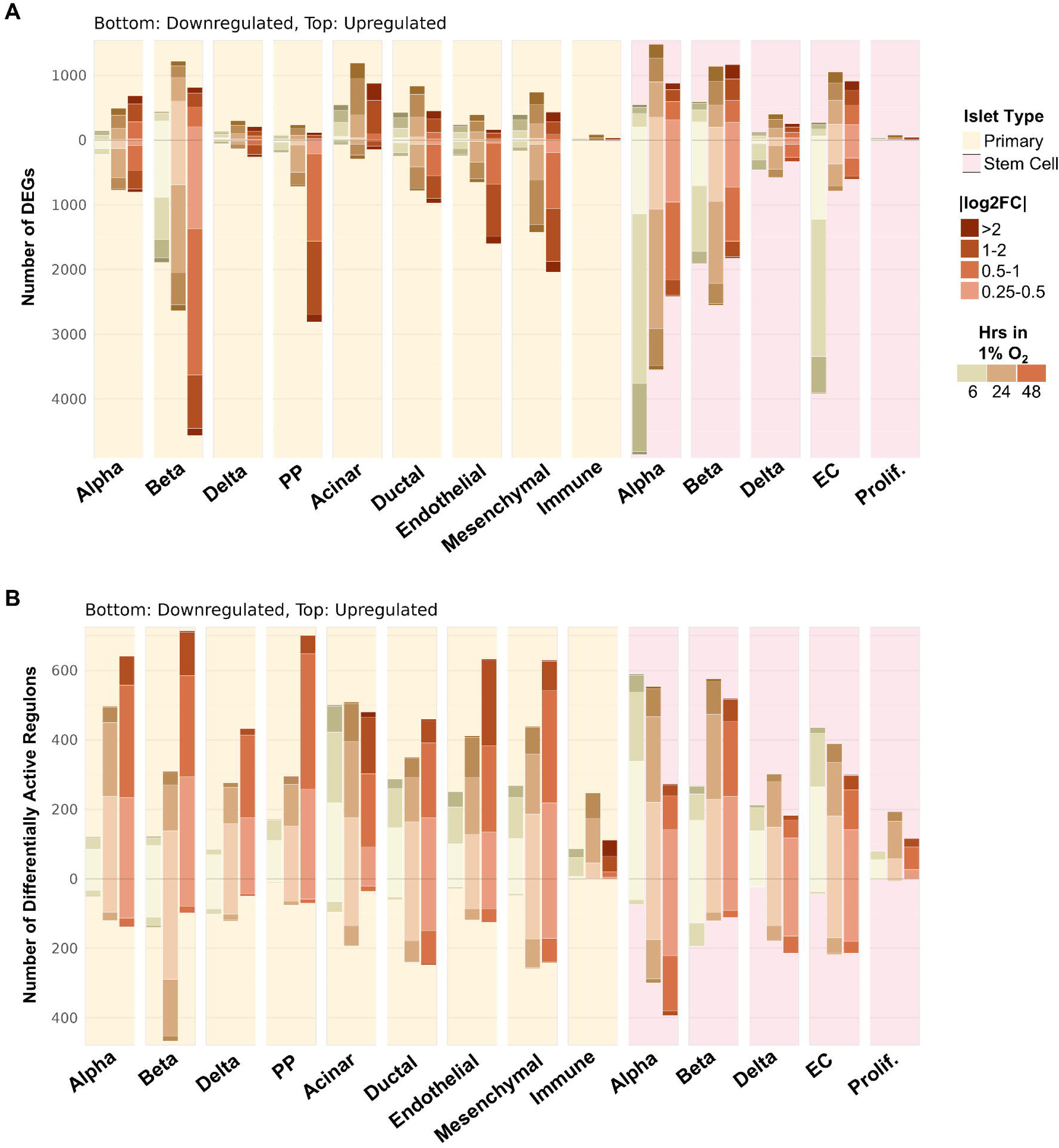
Transcriptional and regulatory dynamics across all identified cell types in response to hypoxia. All comparisons are relative to the 0-hour normoxic baseline for each respective cell type and source. Primary islet cell types are shown with a yellow background; SC-islet cell types are shown with a pink background. (A) Stacked bar charts quantifying the number of differentially expressed genes (DEGs) for each identified cell type at 6, 24, and 48 hours of hypoxia. Bars represent the total number of upregulated (top) and downregulated (bottom) genes. The color shading within each bar indicates the magnitude of the average log_2_ fold-change. (B) Stacked bar charts quantifying the number of differentially active regulons for each cell type, displayed in the same format as (A).

## REFERENCES

1. DiMeglio LA, Evans-Molina C, Oram RA. Type 1 diabetes. Lancet (London, England). 2018 Jun 6;391(10138):2449. Available from: https://www.ncbi.nlm.nih.gov/pmc/articles/PMC6661119/

2. Gong B, Yang W, Xing Y, Lai Y, Shan Z. Global, regional, and national burden of type 1 diabetes in adolescents and young adults. Pediatr Res 2024. 2024 Mar 5;1–9. Available from: https://www.nature.com/articles/s41390-024-03107-5

3. Quattrin T, Mastrandrea LD, Walker LSK. Type 1 diabetes. Lancet. 2023 Jun 24;401(10394):2149–62. Available from: http://www.thelancet.com/article/S0140673623002234/fulltext

4. Holt RIG, Devries JH, Hess-Fischl A, Hirsch IB, Kirkman MS, Klupa T, et al. The Management of Type 1 Diabetes in Adults. A Consensus Report by the American Diabetes Association (ADA) and the European Association for the Study of Diabetes (EASD). Diabetes Care. 2021 Nov 1;44(11):2589–625. Available from: 10.2337/dci21-0043

5. Shapiro AMJ, Lakey JRT, Ryan EA, Korbutt GS, Toth E, Warnock GL, et al. Islet Transplantation in Seven Patients with Type 1 Diabetes Mellitus Using a Glucocorticoid-Free Immunosuppressive Regimen. N Engl J Med. 2000 Jul 27;343(4):230–8. Available from: https://www.nejm.org/doi/full/10.1056/NEJM200007273430401

6. Scharp DW, Lacy PE, Santiago J V., Mccullough CS, Weide LG, Falqui L, et al. Insulin Independence After Islet Transplantation Into Type I Diabetic Patient. Diabetes. 1990 Apr 1;39(4):515–8. Available from: 10.2337/diab.39.4.515

7. Wong JM, Pepper AR. Status of islet transplantation and innovations to sustainable outcomes: novel sites, cell sources, and drug delivery strategies. Front Transplant. 2024 Nov 1;3:1485444.

8. Pagliuca FW, Millman JR, Gürtler M, Segel M, Van Dervort A, Ryu JH, et al. Generation of Functional Human Pancreatic β Cells In Vitro. Cell. 2014 Oct 9;159(2):428–39. Available from: https://pubmed.ncbi.nlm.nih.gov/25303535/

9. Rezania A, Bruin JE, Arora P, Rubin A, Batushansky I, Asadi A, et al. Reversal of diabetes with insulin-producing cells derived in vitro from human pluripotent stem cells. Nat Biotechnol. 2014 Nov 11;32(11):1121–33. Available from: http://www.nature.com/articles/nbt.3033

10. Millman JR, Xie C, Van Dervort A, Gürtler M, Pagliuca FW, Melton DA. Generation of stem cell-derived β-cells from patients with type 1 diabetes. Nat Commun 2016 71. 2016 May 10;7(1):1–9. Available from: https://www.nature.com/articles/ncomms11463

11. Vegas AJ, Veiseh O, Gürtler M, Millman JR, Pagliuca FW, Bader AR, et al. Long-term glycemic control using polymer-encapsulated human stem cell–derived beta cells in immune-competent mice. Nat Med 2015 223. 2016 Jan 25;22(3):306–11. Available from: https://www.nature.com/articles/nm.4030

12. D’Amour KA, Bang AG, Eliazer S, Kelly OG, Agulnick AD, Smart NG, et al. Production of pancreatic hormone–expressing endocrine cells from human embryonic stem cells. Nat Biotechnol. 2006 Oct 19;24(11):1392–401. Available from: https://www.nature.com/articles/nbt1259

13. Millman JR, Pagliuca FW. Autologous Pluripotent Stem Cell–Derived β-Like Cells for Diabetes Cellular Therapy. Diabetes. 2017 May 1;66(5):1111–20. Available from: 10.2337/db16-1406

14. Hogrebe NJ, Maxwell KG, Augsornworawat P, Millman JR. Generation of insulin-producing pancreatic β cells from multiple human stem cell lines. Nat Protoc. 2021;16(9):4109–43.

15. Barsby T, Ibrahim H, Lithovius V, Montaser H, Balboa D, Vähäkangas E, et al. Differentiating functional human islet-like aggregates from pluripotent stem cells. STAR Protoc. 2022 Dec 12;3(4). Available from: /pmc/articles/PMC9508476/

16. Lithovius V, Lahdenpohja S, Ibrahim H, Saarimäki-Vire J, Uusitalo L, Montaser H, et al. Non-invasive quantification of stem cell-derived islet graft size and composition. Diabetologia. 2024 Sep 1;67(9):1912. Available from: https://pmc.ncbi.nlm.nih.gov/articles/PMC11410899/

17. Balboa D, Barsby T, Lithovius V, Saarimäki-Vire J, Omar-Hmeadi M, Dyachok O, et al. Functional, metabolic and transcriptional maturation of human pancreatic islets derived from stem cells. Nat Biotechnol. 2022 Jul 1;40(7):1042. Available from: https://pmc.ncbi.nlm.nih.gov/articles/PMC9287162/

18. Velazco-Cruz L, Song J, Maxwell KG, Goedegebuure MM, Augsornworawat P, Hogrebe NJ, et al. Acquisition of Dynamic Function in Human Stem Cell-Derived β Cells. Stem Cell Reports. 2019 Feb 12;12(2):351. Available from: https://pmc.ncbi.nlm.nih.gov/articles/PMC6372986/

19. Iworima DG, Rieck S, Kieffer TJ. Process parameter development for the scaled generation of stem cell derived pancreatic endocrine cells. Stem Cells Transl Med. 2021 Nov 1;10(11):1459. Available from: https://pmc.ncbi.nlm.nih.gov/articles/PMC8550703/

20. Hering BJ, Rickels MR, Bellin MD, Millman JR, Tomei AA, García AJ, et al. Advances in Cell Replacement Therapies for Diabetes. Diabetes. 2025 Apr 24; Available from: http://www.ncbi.nlm.nih.gov/pubmed/40272266

21. Reichman TW, Markmann JF, Odorico J, Witkowski P, Fung JJ, Wijkstrom M, et al. Stem Cell–Derived, Fully Differentiated Islets for Type 1 Diabetes. N Engl J Med. 2025 Jun 20; Available from: https://pubmed.ncbi.nlm.nih.gov/40544428/

22. Komatsu H, Kandeel F, Mullen Y. Impact of Oxygen on Pancreatic Islet Survival. Pancreas. 2018 May 1;47(5):533–43. Available from: https://journals.lww.com/00006676-201805000-00004

23. Komatsu H, Cook C, Wang CH, Medrano L, Lin H, Kandeel F, et al. Oxygen environment and islet size are the primary limiting factors of isolated pancreatic islet survival. PLoS One. 2017 Aug 1;12(8):1–17. Available from: https://journals.plos.org/plosone/article?id=10.1371/journal.pone.0183780

24. Olsson R, Olerud J, Pettersson U, Carlsson PO. Increased numbers of low-oxygenated pancreatic islets after intraportal islet transplantation. Diabetes. 2011 Sep;60(9):2350–3. Available from: https://pmc.ncbi.nlm.nih.gov/articles/PMC3161309/

25. Carlsson PO, Liss P, Andersson A, Jansson L. Measurements of oxygen tension in native and transplanted rat pancreatic islets. Diabetes. 1998 Jul 1;47(7):1027–32. Available from: 10.2337/diabetes.47.7.1027

26. Tsuyama T, Sato Y, Yoshizawa T, Matsuoka T, Yamagata K. Hypoxia causes pancreatic β cell dysfunction and impairs insulin secretion by activating the transcriptional repressor BHLHE40. 2023 Aug 3;24(8):e56227. Available from: https://pmc.ncbi.nlm.nih.gov/articles/PMC10398664/

27. Delaune V, Berney T, Lacotte S, Toso C. Intraportal islet transplantation: the impact of the liver microenvironment. Transpl Int. 2017 Mar 1;30(3):227–38. Available from: https://onlinelibrary.wiley.com/doi/full/10.1111/tri.12919

28. Rodriguez-Brotons A, Bietiger W, Peronet C, Magisson J, Sookhareea C, Langlois A, et al. Impact of Pancreatic Rat Islet Density on Cell Survival during Hypoxia. J Diabetes Res. 2015;2016:3615286. Available from: https://pmc.ncbi.nlm.nih.gov/articles/PMC4707363/

29. Carlsson O, Palm F, Andersson A, Liss P, Carlsson PO, Palm F, et al. Markedly Decreased Oxygen Tension in Transplanted Rat Pancreatic Islets Irrespective of the Implantation Site [Internet]. Vol. 50, Diabetes. American Diabetes Association; 2001 Mar. Available from: 10.2337/diabetes.50.3.489

30. Liljebäck H, Grapensparr L, Olerud J, Carlsson POO. Extensive Loss of Islet Mass beyond the First Day after Intraportal Human Islet Transplantation in a Mouse Model. Cell Transplant. 2016 Mar 1;25(3):481–9. Available from: https://journals.sagepub.com/doi/10.3727/096368915X688902

31. Faleo G, Russ HA, Wisel S, Parent A V, Nguyen V, Nair GG, et al. Mitigating Ischemic Injury of Stem Cell-Derived Insulin-Producing Cells after Transplant. Stem cell reports. 2017 Sep 12;9(3):807–19. Available from: http://www.ncbi.nlm.nih.gov/pubmed/28803916

32. Biarnés M, Montolio M, Nacher V, Raurell M, Soler J, Montanya E. β-Cell Death and Mass in Syngeneically Transplanted Islets Exposed to Short-and Long-Term Hyperglycemia. Diabetes. 2002 Jan 1;51(1):66–72. Available from: 10.2337/diabetes.51.1.66

33. Davalli AM, Scaglia L, Zangen DH, Hollister J, Bonner-Weir S, Weir GC. Vulnerability of Islets in the Immediate Posttransplantation Period: Dynamic Changes in Structure and Function. Diabetes. 1996 Sep 1;45(9):1161–7. Available from: http://diabetesjournals.org/diabetes/article-pdf/45/9/1161/555374/45-9-1161.pdf

34. Lehmann R, Zuellig RA, Kugelmeier P, Baenninger PB, Moritz W, Perren A, et al. Superiority of Small Islets in Human Islet Transplantation. Diabetes. 2007 Mar 1;56(3):594–603. Available from: https://diabetesjournals.org/diabetes/article/56/3/594/15115/Superiority-of-Small-Islets-in-Human-Islet

35. Buchwald P. FEM-based oxygen consumption and cell viability models for avascular pancreatic islets. Theor Biol Med Model. 2009 Apr 16;6(1):1–13. Available from: https://tbiomed.biomedcentral.com/articles/10.1186/1742-4682-6-5

36. Bensellam M, Maxwell EL, Chan JY, Luzuriaga J, West PK, Jonas JC, et al. Hypoxia reduces ER-to-Golgi protein trafficking and increases cell death by inhibiting the adaptive unfolded protein response in mouse beta cells. Diabetologia. 2016 Jul 4;59(7):1492–502. Available from: http://link.springer.com/10.1007/s00125-016-3947-y

37. Narimiya M, Yamada H, Matsuba I, Ikeda Y, Tanese T, Abe M. The Effect of Hypoxia on Insulin and Glucagon Secretion in the Perfused Pancreas of the Rat. Endocrinology. 1982 Sep 1;111(3):1010–4. Available from: 10.1210/endo-111-3-1010

38. Dionne KE, Colton CK, Yarmush ML. Effect of Hypoxia on Insulin Secretion by Isolated Rat and Canine Islets of Langerhans. Diabetes. 1993 Jan 1;42(1):12–21. Available from: 10.2337/diab.42.1.12

39. Sato Y, Inoue M, Yoshizawa T, Yamagata K. Moderate Hypoxia Induces β-Cell Dysfunction with HIF-1–Independent Gene Expression Changes. PLoS One. 2014 Dec 12;9(12):e114868. Available from: https://journals.plos.org/plosone/article?id=10.1371/journal.pone.0114868

40. Kitzmann JP, O’Gorman D, Kin T, Gruessner AC, Senior P, Imes S, et al. Islet Oxygen Consumption Rate Dose Predicts Insulin Independence for First Clinical Islet Allotransplants. Transplant Proc. 2014 Jul 1;46(6):1985–8.

41. Davis JC, Alves TC, Helman A, Chen JC, Kenty JH, Cardone RL, et al. Glucose Response by Stem Cell-Derived β Cells In Vitro Is Inhibited by a Bottleneck in Glycolysis. Cell Rep. 2020 May 12;31(6):107623. Available from: https://pmc.ncbi.nlm.nih.gov/articles/PMC7433758/

42. Thorngren J, Brboric A, Vasylovska S, Hjelmqvist D, Westermark GT, Saarimäki-Vire J, et al. Efficient Vascular and Neural Engraftment of Stem Cell–Derived Islets. Diabetes. 2024 Jul 1;73(7):1127. Available from: https://pmc.ncbi.nlm.nih.gov/articles/PMC11189832/

43. Stokes RA, Cheng K, Deters N, Lau SM, Hawthorne WJ, O’connell PJ, et al. Hypoxia-Inducible Factor-1α (HIF-1α) Potentiates β-Cell Survival after Islet Transplantation of Human and Mouse Islets. Cell Transplant. 2013 Feb 1;22(2):253–66. Available from: https://pmc.ncbi.nlm.nih.gov/articles/PMC6595221/

44. Cheng K, Ho K, Stokes R, Scott C, Lau SM, Hawthorne WJ, et al. Hypoxia-inducible factor-1α regulates β cell function in mouse and human islets. J Clin Invest. 2010 Jun 1;120(6):2171–83. Available from: http://www.jci.org/articles/view/35846

45. Cantley J, Grey ST, Maxwell PH, Withers DJ. The hypoxia response pathway and β-cell function. Diabetes, Obes Metab. 2010;12(SUPPL. 2):159–67. Available from: https://onlinelibrary.wiley.com/doi/full/10.1111/j.1463-1326.2010.01276.x

46. Yamagata K, Tsuyama T, Sato Y. Roles of β-Cell Hypoxia in the Progression of Type 2 Diabetes. Int J Mol Sci. 2024 Apr 1;25(8):4186. Available from: https://pmc.ncbi.nlm.nih.gov/articles/PMC11050445/

47. Augsornworawat P, Hogrebe NJ, Ishahak M, Schmidt MD, Marquez E, Maestas MM, et al. Single-nucleus multi-omics of human stem cell-derived islets identifies deficiencies in lineage specification. Nat Cell Biol. 2023 Jun 1;25(6):904. Available from: /pmc/articles/PMC10264244/

48. Maestas MM, Ishahak M, Augsornworawat P, Veronese-Paniagua DA, Maxwell KG, Velazco-Cruz L, et al. Identification of unique cell type responses in pancreatic islets to stress. Nat Commun 2024 151. 2024 Jul 2;15(1):1–17. Available from: https://www.nature.com/articles/s41467-024-49724-w

49. Schmidt MD, Ishahak M, Augsornworawat P, Millman JR. Comparative and integrative single cell analysis reveals new insights into the transcriptional immaturity of stem cell-derived β cells. BMC Genomics. 2024 Dec 1;25(1). Available from: https://pubmed.ncbi.nlm.nih.gov/38267908/

50. Veres A, Faust AL, Bushnell HL, Engquist EN, Kenty JHR, Harb G, et al. Charting cellular identity during human in vitro β-cell differentiation. Nature. 2019 May 16;569(7756):368–73. Available from: http://www.ncbi.nlm.nih.gov/pubmed/31068696

51. Weng C, Xi J, Li H, Cui J, Gu A, Lai S, et al. Single-cell lineage analysis reveals extensive multimodal transcriptional control during directed beta-cell differentiation. Nat Metab 2020 212. 2020 Nov 30;2(12):1443–58. Available from: https://www.nature.com/articles/s42255-020-00314-2

52. Zhu H, Wang G, Nguyen-Ngoc KV, Kim D, Miller M, Goss G, et al. Understanding cell fate acquisition in stem-cell-derived pancreatic islets using single-cell multiome-inferred regulomes. Dev Cell. 2023 May 8;58(9):727–743.e11.

53. Wang X, Brielle S, Kenty-Ryu J, Korover N, Bavli D, Pop R, et al. Improving cellular fitness of human stem cell-derived islets under hypoxia. Nat Commun 2025 161. 2025 May 23;16(1):1–17. Available from: https://www.nature.com/articles/s41467-025-59924-7

